# Microscopy quantification of microbial birth and death dynamics

**DOI:** 10.1101/324269

**Authors:** Samuel F. M. Hart, David Skelding, Adam J. Waite, Justin Burton, Li Xie, Wenying Shou

## Abstract

Microbes live in dynamic environments where nutrient concentrations fluctuate. Quantifying fitness (birth and death) in a wide range of environments is critical for understanding microbial evolution as well as ecological interactions where one species alters the fitness of another. Here, using high-throughput time-lapse microscopy, we have quantified how *Saccharomyces cerevisiae* mutants incapable of synthesizing an essential metabolite grow or die in various concentrations of the required metabolite. We establish that cells normally expressing fluorescent proteins lose fluorescence upon death and that the total fluorescence in an imaging frame is proportional to the number of live cells even when cells form multiple layers. We validate our microscopy approach of measuring birth and death rates using flow cytometry, cell counting, and chemostat culturing. For lysine-requiring cells, very low concentrations of lysine are not detectably consumed and do not support cell birth, but delay the onset of death phase and reduce the death rate. In contrast, in low hypoxanthine, hypoxanthine-requiring cells can produce new cells, yet also die faster than in the absence of hypoxanthine. For both strains, birth rates under various metabolite concentrations are better described by the sigmoidal-shaped Moser model than the well-known Monod model, while death rates depend on the metabolite concentration and can vary with time. Our work reveals how time-lapse microscopy can be used to discover non-intuitive microbial dynamics and to quantify growth rates in many environments.

## Introduction

Understanding microbial evolution and ecology requires quantifying microbial fitness in diverse environments that the microbes typically encounter. Fitness is often measured as the net growth rate (“growth rate”) – the difference between birth and death rates.

An easy and rapid method for measuring growth rate is to track optical density of a culture over time. This method is useful when death rate is low, since optical density cannot differentiate between live and dead cells. In contrast, flow cytometry can yield live and dead cell counts, but requires periodic manual sampling of the culture under observation. As an alternative method, high-throughput microscopy has been developed and applied to, for example, monitoring biofilm susceptibility to antibiotics ^1^, quantifying growth rate heterogeneity among microcolonies ^2^, and screening large collections of mutant strains ^3^.

Here, we use microscopy to distinguish cell birth from cell death, especially at low metabolite concentrations where death rate is high. Distinguishing birth from death can be important. For example, metabolite consumption is tied to birth and not to death. As another example, in the extreme case of cells not dividing or dying, then natural selection ceases. Based on the same reasoning, if two populations have the same net growth rate, then a population that divides and dies slowly should evolve slower per unit time compared to a population that divides and dies rapidly.

Several mathematical models phenomenologically relate nutrient concentrations to population growth rates. The best known model is the Monod model ^4^, *ɡ* = *ɡ*_*max*_ *s*/(*K*_*M*_ + *s*), where *g* is the net growth rate, *s* is the concentration of the limiting metabolite, *g*_*max*_ is the maximal growth rate, and *K*_*M*_ is the concentration of *s* at which half *g*_*max*_ is achieved. Other growth models such as the Teisser and the Contois ^5^ models have also been proposed. However, like the Monod model, they do not consider cell death since they assume zero (instead of negative) growth rate at zero metabolite concentration. A different growth model by Kovárová-Kovar and Egli ^6^ incorporates a fixed death rate, although in reality, death rate could vary with metabolite concentrations.

Previously, we constructed a two-strain synthetic yeast cooperative community as a model system to explore how cells in a cooperative community might evolve and how cooperation might shape species coexistence and spatial patterning ^7–11^. In this community, a red-fluorescent strain required lysine and released hypoxanthine (an adenine precursor) (BioRxiv), while a green-fluorescent strain required hypoxanthine and released lysine. A mathematical model for this community had model parameters including each strain’s birth and death rates at various concentrations of the required metabolite, metabolite release rate, and metabolite consumption per birth.

Here, we describe a high-throughput microscopy assay that we developed and validated for quantifying a strain’s birth and death rates at various concentrations of the required metabolite. Our approach can be applied to quantifying the birth and death dynamics of other fluorescently labeled microbes.

## Results

### Using fluorescence to quantify cell birth and death

Our inverted fluorescence microscope is equipped with motorized stage and filter wheel, and is enclosed in a temperature-controlled chamber (Figure 1A) to ensure a nearly constant temperature (Supp Fig 1; Supp Fig 10). To enable automated long-term imaging with minimal photo-damage, we wrote a LabView routine to perform autofocusing using the bright field, and then imaged in the fluorescence channel. However, despite controlling the temperature, condensation developed on the microplate lid over time,
 which sometimes interfered with autofocusing. To resolve this, we developed a “lid warmer” using transparent, conductive ITO glass (Figure 1B) to warm the microplate lid to ~0.7°C above the stage temperature (Supp Fig 1). This eliminated condensation (Figure 1C) and allowed reliable auto-focusing over tens of hours.

**Figure 1.**
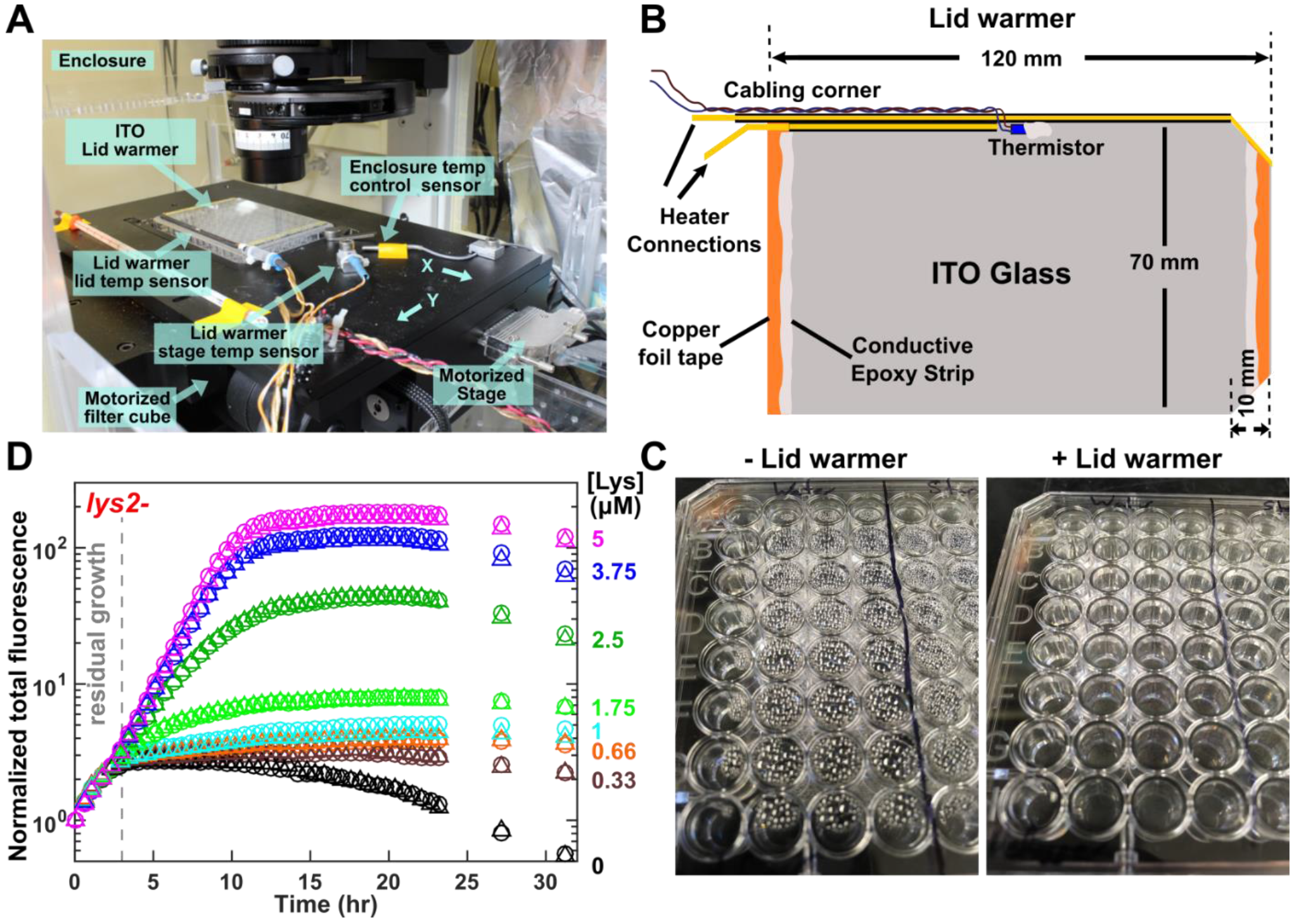
Automated high-throughput microscopy. **(A)** Microscope setup. An enclosure around the microscope provides a temperature-controlled environment. A motorized stage allows repeated bright field and fluorescent imaging of the same positions in specified wells of a microtiter plate (Supp Movies 1-3). Motorized filter cubes allows automated filter switching. **(B)** An ITO glass lid warmer prevents condensation. Sensors on the plate lid (“thermistor”) and the microscope stage provide temperature measurements to a LabVIEW program which turns the ITO lid warmer on or off to maintain the plate at ~ 0.7°C warmer than the stage (Supp Fig 1). **(C)** The lid warmer eliminates condensation. The images were taken after a 24-hour imaging experiment at 30°C. **(D)** Growth of *lys2*- (WY1335) cells at various lysine concentrations. Background-subtracted total fluorescence intensities from four picture frames were normalized against their respective initial values, averaged, and plotted. Dashed line marks the end of residual growth. Residual growth occurs even at zero lysine, and is presumably fueled by vacuolar lysine storage. When calculating growth rates, we only considered post residual growth data. Circles and triangles mark two independent experiments.

For yeast cells expressing a fluorescent protein, total fluorescence intensity (after background subtraction) scaled linearly with live fluorescent cell density up to at least nine cell layers (Supp Fig 2). Occasionally, we also observed that cells lost fluorescence immediately upon losing cell integrity (compare Supp Movie 3 vs Supp Movie 5). Thus, increases or decreases in fluorescence were proportional to cell division or cell death, respectively.

To measure birth and death rates, we performed time-lapse imaging of fluorescent yeast cells at various concentrations of the required metabolite (e.g. Figure 1D). We started with a small number of cells to minimize metabolite depletion during growth rate measurements. Even in the absence of the required metabolite, total fluorescence intensity initially increased due to residual growth fueled by cellular storage of metabolites ^12^ (e.g. 0~3 hrs in Figure 1D). Thus, we only used images after the residual growth in our data analysis.

### Death rate is time-dependent

Death rate is not a constant. For example, we measured the death rate of *lys2*- cells in zero lysine. Since new birth was negligible (no birth out of 603 cells over 30 hrs; Supp Movie 3), death rate could be estimated by quantifying the negative slope of ln(live population size) against time. We observed multi-phasic death kinetics, with a slow death rate followed by a faster death rate (Figure 2A, lightest grey; Figure 2B, [Lys]=0; Supp Fig 3). *ade8*- cells in the absence of hypoxanthine also displayed a time-dependent death rate (Supp Fig 4). For both strains, the death rate would eventually slow down, as shown in our previous work ^7^. As we will demonstrate below, death rate also depends on the metabolite concentration.

**Figure 2.**
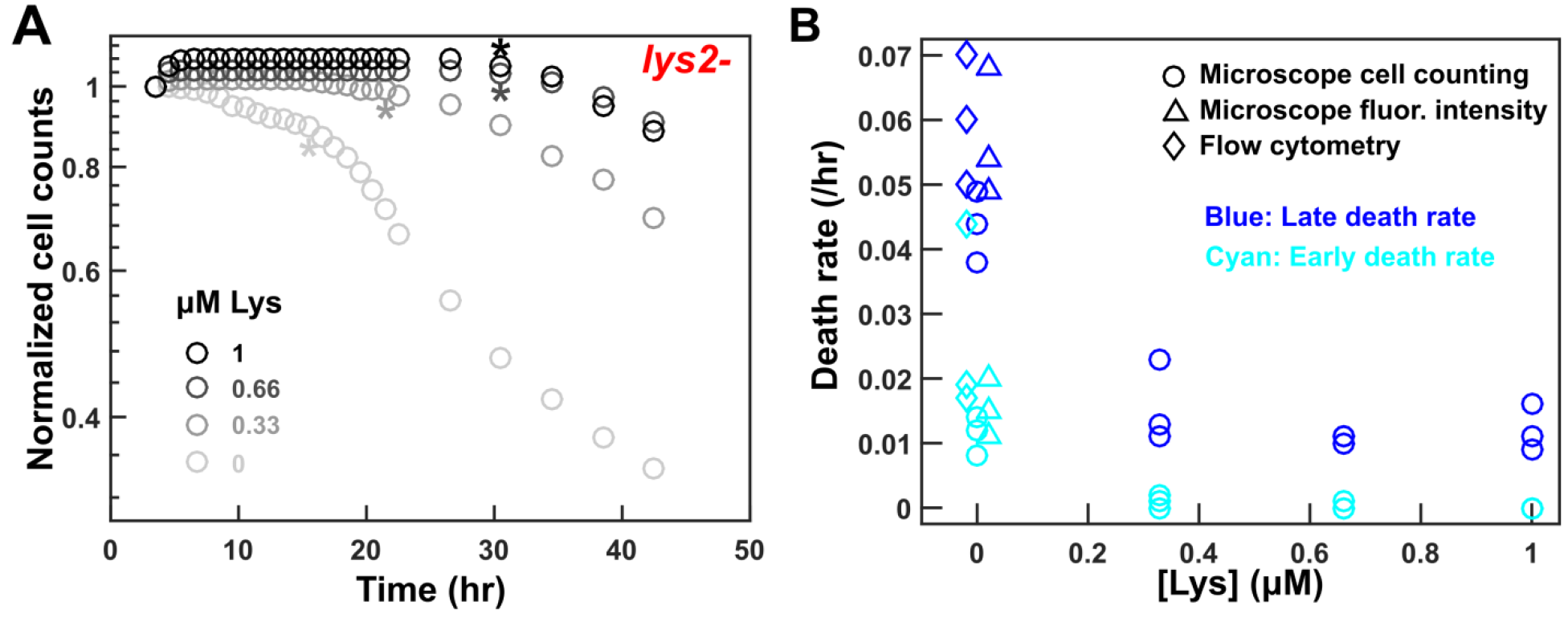
Death rate is nutrient and time-dependent. Exponentially-growing *lys2*- (WY1335) cells were washed free of lysine at 0 hr, and imaged in minimal medium supplemented with 0 to 1 μM lysine. **(A)** Birth events were restricted to the first few hours, and may be regarded as part of residual growth. Early and late death rates were calculated using data before and after asterix (*), respectively. For each experiment, we tracked birth and death events from an initial 200~300 cells. **(B)** Death rate started slow (cyan), and then increased (blue). For experiments in 0 μM lysine, data points are jittered along the x-axis to aid visualization. Death rates of *lys2*- cells at 0 μM lysine were tracked using microscopy cell counting (circles), microscopy fluorescence intensity (triangles), and flow cytometry (diamonds). The three methods resulted in compatible results, although death rate from cell counting seemed to be lower than that from microscopy fluorescence intensity or flow cytometry. Death rates at 0.33~1 μM lysine were tracked using microscopy cell counting only (circles).

Death rate based on total fluorescence is comparable to two other approaches where fluorescent cells were scored “live” and non-fluorescent cells were scored “dead”: flow cytometry and direct cell counting in microscopy images. As expected, subtle differences existed among the three methods (Supp Fig 3). For example, total fluorescence intensity (but not flow cytometry or cell counting) would be increased by cell swelling (Figure 3C). The microscopy cell counting method could detect the death of a bud, but in flow cytometry, since the attached mother cell was still fluorescent, the death event would not be recorded (unless a death dye was used). Overall, results from microscopy fluorescence and flow cytometry overlapped, while that from cell counting yielded a slightly lower death rate (Figure 2B, compare different symbols of the same color), possibly because sample handling in flow cytometry reduced the viability of starving cells.

**Figure 3.**
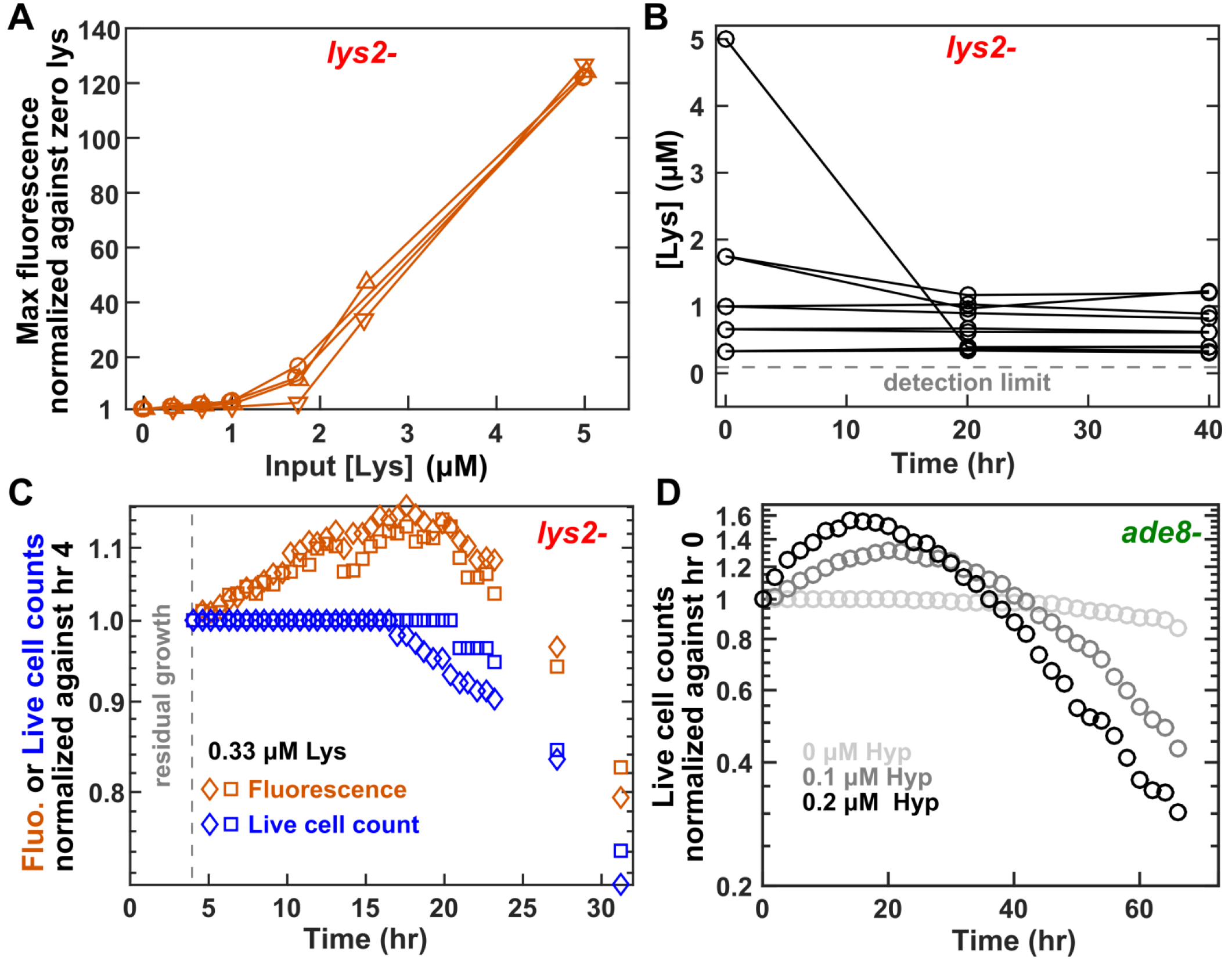
Very low concentrations of metabolites may not be consumed and can lead to diverse birth and death dynamics. **(A)** Maximal fluorescence achieved by a *lys2*- (WY1335) culture scales with initial lysine concentration only for lysine concentrations above a threshold (1.75 μM). Maximal fluorescence was normalized against that at zero lysine. **(B)** Low concentrations of lysine (≤1 μM) remain largely unconsumed by *lys2*- over 40 hours. We measured lysine concentrations in supernatants using the rate bioassay (Methods). **(C)** An increase in fluorescence intensity may not correspond to cell birth. We imaged *lys2*- cells (161 total) in 0.33 μM lysine. Total fluorescence intensity (brown) and counts of fluorescent cells (blue) over time are plotted. We observed no birth events in this experiment, while fluorescence increased for over 15 hours after the start of the experiment. **(D)** A low level of hypoxanthine increases cell birth and death in *ade8*- cells. Exponential *ade8*- (WY1340) cells were washed and starved for 24 hours, and then imaged every 2 hours in 0 (light grey), 0.1 μM (dark grey) and 0.2 μM (black) hypoxanthine. A small amount of hypoxanthine increases both cell birth (0~20 hours; Supp Fig 5B) and cell death (20-66 hours).

### Diverse birth and death dynamics at low metabolite concentrations

Low concentrations of metabolites may not be consumed. For example, low concentrations of lysine (e.g. 0.33~1 μM; Supp Movie 2) was barely consumed by *lys2*- cells, while concentrations >1.75 μM were depleted down to <1 μM (Fig 3B). Consistently, maximal fluorescence intensity only scaled linearly with lysine concentrations > 1.75 μM (Figure 3A), the level around which half maximal growth rate was achieved (Figure 4). For *ade8*- cells at low input concentrations (e.g. 0.1~0.2 μM) of hypoxanthine, supernatant hypoxanthine concentration was too low to be directly measured. However, we inferred that low concentrations of hypoxanthine were largely un-consumed based on the following inference. Since 1~3 fmole of hypoxanthine is consumed per *ade8*- cell (BioRxiv), the input medium (300 μl of 0.1 μM or 3×10^4^ fmole hypoxanthine) should support the birth of a total of 1×10^4^~3×10^4^ cells. Instead, starting with 3000 cells, we observed <50% increase in cell number (Figure 3D).

Low concentrations of metabolites lead to diverse birth and death dynamics depending on the strain genotype. For *lys2*- cells in low lysine, although total fluorescence intensity increased for longer compared to zero lysine (Figure 1D), this increase generally corresponded to cell swelling rather than cell birth (Figure 3C). Birth events, if any, were restricted to the initial few hours and not sustained at later time points despite nearly-constant metabolite concentration (Figure 2A; Figure 3B; Supp Fig 5A). Thus, the initial birth events could be interpreted as low lysine prolonging the residual growth phase and delaying the onset of death phase. Low lysine also reduced early and late death rates (Figure 2B). In contrast, for *ade8*- cells, low input concentrations (e.g. 0.1~0.2 μM) of hypoxanthine led to increased birth and death rates compared to zero hypoxanthine (Figure 3D). *ade8*- cells were occasionally born in low hypoxanthine, even after the onset of death phase (30-66 hr, Supp Fig 5B). Moreover, a small number of *ade8*- cells transiently lost fluorescence, but then regained fluorescence and continued to divide (Supp Movie 4). In summary, for *lys2*- cells, low concentrations of lysine were not consumed, did not sustain birth beyond the initial stage, but delayed the onset of death phase and slowed the death rate once cells began to die (Figure 2). For *ade8*- cells, low concentrations of hypoxanthine were likely also largely unconsumed, and increased both birth and death rates. Overall, death rate was not only time-dependent, but also nutrient concentration-dependent.

### The Moser model is superior to the Monod model in describing lys2- and ade8- cell birth

At relatively high lysine concentrations (≥1.75 μM, Supp Movie 1), we could not directly count birth and death events due to multiple cell layers. Since the death rate was small (<0.002/hr) compared to the net growth rate (≥ 0.1/hr) in this range of lysine concentrations (BioRxiv), we approximated the net growth rate as the birth rate (Figure 4). That is, we measured the rate of ln(total fluorescence) increase across sliding time windows (generally four time points over a total of 1.7 hrs; Figure 1D), and used the steepest slope as birth rate.

When lysine was not the limiting nutrient in the media (≥ 5 μM), *lys2*- cells achieved a maximal net growth rate of 0.50 +/− 0.02/hr in microscopy assay (Figure 1D, magenta), consistent with the value measured from culture optical density over time (e.g. 0.49 +/− 0.03/hr). For lysine concentrations ≥ 2.5 μM, the maximal growth rate was maintained across at least two contiguous sliding windows (Supp Fig 6). However, at lower lysine concentrations (e.g. 1.75 μM), the growth rate continuously declined throughout the experiment, and thus it was unclear whether our measured maximal birth rate was truly maximal. To verify our observations, we employed an independent measurement method where we grew *lys2*- cells in lysine-limited chemostats ^13^ (Methods). In a steady-state chemostat, the population net growth rate is equal to the dilution rate. We set the dilution rate to various values, and measured the corresponding lysine concentrations (Supp Fig 7). Chemostat measurements were consistent with microscopy measurements (Figure 4, blue crosses).

The birth rate of *lys2*- cell increased with lysine concentration in a sigmoidal fashion. The data were better characterized by the Moser model *b*(*s*) = *b*_*max*_*s*^*n*^ / (*K*_*m*_^*n*^ + *s*^*n*^) (black line; Figure 4) than the Monod model *b*(*s*) = *b*_*max*_*s* / (*K*_*m*_ + *s*)(grey dotted line; Figure 4) In both models, *s* is lysine concentration, *b*_*max*_ is the maximal birth rate, *K*_*m*_ is the Monod constant (*s* required to achieve half maximal birth rate). The Moser model has an additional parameter, *n*, which is analogous to the cooperativity coefficient in the Hill equation.

**Figure 4.**
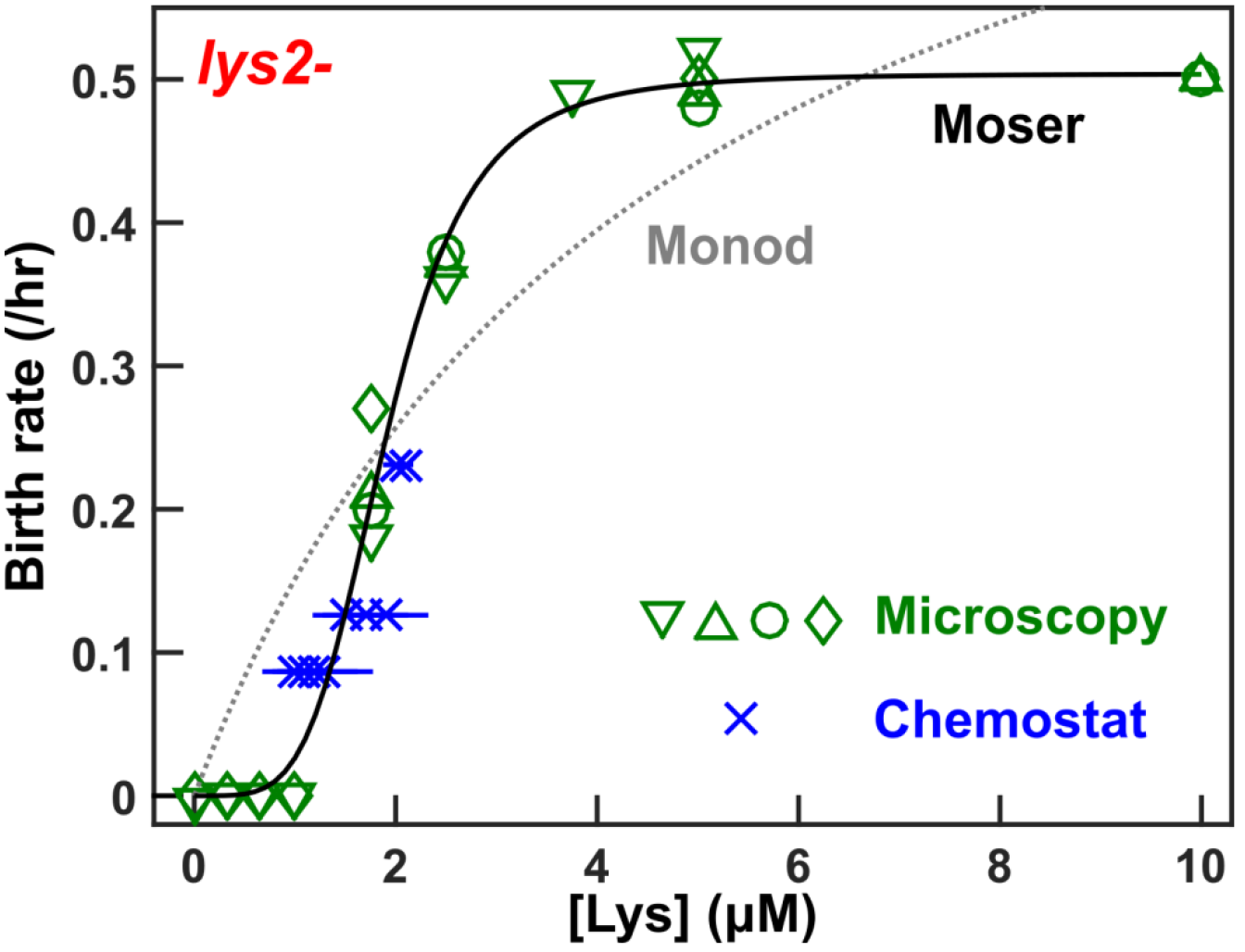
The birth rate of lys2- as a function of lysine concentration is better described by the Moser model than the Monod model. For lysine concentrations ≥1.75 μM, we calculated net growth rates within 3~4 hr sliding windows. We used the maximal growth rates (before lysine depletion) to approximate birth rates, since death rates are at most 2% of growth rates and can thus be neglected (see Figure 2). At lower lysine concentrations (≤1μM), cells remain monolayer, which has enabled us to count birth (and death) events. Four microscopy assays are shown (green) with each assay designated by a different symbol. Despite increasing fluorescence at low lysine concentrations, we observed no births (e.g. Figure 3C). Growth rates and their approximate birth rates were also measured in chemostats (blue) where growth rates are controlled via dilution rates. The corresponding lysine concentrations in culturing vessels were measured using a bioassay (Methods). Results from different measurements are consistent‥ The birth rate of *lys2*- as a function of lysine concentration can be described by the Moser equation (black) where 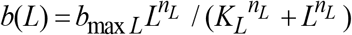 with *b*_*max L*_ = 0.50 (95% CI: 0.48 ~ 0.53), *K*_*L*_ = 1.9 (95% CI: 1.8 ~ 2.0), and *n* = 4.47 (95% CI: 3.2~ 5.7). In comparison, the best-fit of the Monod equation *g*(*L*) =*b*_*maxL*_*L*/(*K*_*L*_ + *L*)(grey dotted, *b*_*max L*_ = 0.85 and *K*_*L*_ = 4.6) fits the data poorly.

For *ade8*-, the Monod model worked relatively well (Supp Fig 8, grey), but the Moser model provided a more accurate estimate of the maximal birth rate (Supp Fig 8, black). Experimentally measured maximal birth rate was 0.44+/− 0.03/hr, comparable to the fit by the Moser model (0.44/hr) and significantly lower than the fit by the Monod model (0.49/hr). Increased accuracy in the Moser model is not surprising given the additional parameter. Nevertheless, an accurate fit to experimental data is useful when modeling population dynamics.

## Discussion

We demonstrated that our microscopy setup allows quantitative measurements of cell growth in a wide range of nutrient environments. Using this approach, we tracked individual birth and death events in a monolayer at low metabolite concentrations. At higher metabolite concentrations, we quantified the net growth rate which was approximately the birth rate since death rate was relatively small.

Microscopy quantification of growth rates requires careful cell preparation and image analysis. Growth rates can differ depending on the growth conditions before starvation (Supp Fig 9), how we define residual growth (Supp Fig 6), and whether birth and death rates are calculated from fluorescence or cell counts (Figure 3C). Since there is no single “correct” procedure, it is important to be aware of the limitations of each method. We recommend cross-checking one method against another, independent, method. For example, we cross-checked microscopy measurements against chemostat measurements when estimating growth-nutrient relationship (Figure 4), and cross-checked microscopy measurements against flow cytometry when estimating the death rate (Supp Fig 3).

Population dynamics of the two strains shared certain features under nutrient limitation. For example, both *lys2*- and *ade8*- cells displayed time and nutrient-dependent death rates (Figure 2; Supp Fig 3; Supp Fig 4). The two strains differed in other aspects. For *lys2*- cells, low concentrations (0.33-1 μM) of lysine were largely un-consumed (Figure 3A and B), and did not support birth beyond residual growth (Supp Fig 5A), but did delay and slow death when compared to full starvation (Figure 2). In contrast, for *ade8*- cells, low concentrations (0.1-0.2 μM) of hypoxanthine increased death rates (Figure 3D) compared to full starvation, and, at the same time, supported some birth (Supp Fig 5B). These seemingly counter-intuitive behaviors probably resulted from the fact that these mutations disrupted biosynthetic pathways that would normally produce the necessary metabolite. Since the cell does not “know” that it cannot make the metabolite, it continues to try to grow and divide in its absence, which results in an abnormally high death rate ^14^.

The Monod model has been observed to fit, for example, the growth rates of yeast strains at various glucose concentrations ^2^. For *lys2*- cells, the Moser model was much better than the Monod model in describing birth rates as a function of lysine concentrations (Figure 4). The Monod model (Figure 4, grey) overestimated the birth rate at low lysine (<2 μM), underestimated the growth rate at medium lysine (2~6 μM), and overestimated the growth rate at high lysine (>6 μM). Experimentally, the birth rate of *lys2*- increased with lysine concentration in a sigmoidal fashion, corresponding to a lack of birth at low lysine (≤1 μM), a sharp increase in birth rate at medium lysine (1.75~2.5 μM), and leveling off at max birth rate at higher lysine (≥3.75 μM). The sigmoidal growth-nutrient relationship could be explained by, for example, cooperative binding by nutrient transporters, as seen in a variety of cells types, including yeast ^15–17^. For *ade8*- cells in hypoxanthine, the Monod model closely fit experimental data. The Moser model was still more accurate than Monod model (Supp Fig 8), although a two-parameter model (Moser) is expected to improve the fit over a one-parameter model (Monod). Regardless, a growth model that faithfully captures experimental observations is useful. For example, when modeling a community of two cross-feeding strains, a Moser model of how fast each strain grows in various concentrations of partner-supplied metabolite can be incorporated into the community dynamics model. In summary, our work demonstrates the potential of microscopy assays in quantifying microbial birth and death dynamics.

## Methods

### Strains and growth medium

We used strains from the RM11 background with the following genetic modifications introduced via transformation. Strain WY1335 (“*lys2*-”) has the genotype of *ho::loxP AMN1-BYste3::Hph fba1::FBA1-mCherry-loxP ade4::ADE4-PUR6 (o/e) lys2::loxP.* Strain WY1340 (“*ade8*-)” has the genotype of *ho::loxP AMN1-BYste3::NATfba1::FBA1-EGFP-loxP lys21::LYS21(o/e) ade8::loxP*. For our bioassay of low metabolite concentrations, we used an evolved clone (WY2270) isolated after *lys2*- had grown for tens of generations under lysine limitation. This clone displayed an increased affinity for lysine. We stored these strains at −80°C in YPD+15% glycerol.

We used rich medium YPD (10 g/L yeast extract, 20 g/L peptone, 20 g/L glucose) for streaking out single colonies and for growing saturated YPD overnight cultures which were then used as inoculum to grow exponential cultures. We found *ade8*- cells could grow to a higher density in YPD if supplemented with 100 μM hypoxanthine. We sterilized YPD media by autoclaving. YPD overnight cultures were stored at room temperature for no more than 4~5 days prior to experiments. We used defined minimal medium SD (6.7 g/L Difco^™^ yeast nitrogen base w/o amino acids, 20 g/L glucose) for all experiments ^18^, with supplemental metabolites added as noted ^19^ To achieve higher reproducibility than autoclaving, we sterilized SD media by filtering through 0.22 μm filters.

We performed all culturing at 29.5±1°C. *lys2*- cells were pre-grown to exponential phase in SD supplemented with excess (164 μM) lysine and washed 3~5 times with SD. Where noted, we starved *lys2*- cells for 3~6 hours to deplete intracellular lysine storage. Otherwise, we did not starve *lys2*- prior to starting an experiment. *ade8*- cells were pre-grown to exponential phase in SD supplemented with excess hypoxanthine (100 μM), washed 3~5 times with SD, and prestarved in SD for 24 hours to deplete cellular storage.

### Microscope setup

Imaging was performed using a Nikon Eclipse TE-2000U inverted fluorescence microscope. A temperature-controlled enclosure (In Vivo Scientific controller, model 300353) maintained the microscope at a temperature close to a set temperature (29.5°C). We noticed that the air conditioning system could cause fluctuations in temperature, so we turned it off.

The microscope had a motorized stage to allow z-autofocusing (Methods, “Autofocusing“) and systematic xy-scanning of locations in microplate wells. Our LabVIEW (National Instruments) program controlled the microscope body, illumination, and stage position through serial port communication. The program moved the stage gently, so that cells were not disturbed and individual cells could be tracked from one time point to the next. The microscope was also equipped with motorized switchable filter cubes capable of detecting a variety of fluorophores. We used an ET DsRed filter cube (Exciter: ET545/30x, Emitter: ET620/60m, Dichroic: T570LP) for mCherry-expressing strains, and an ET GFP filter cube (Exciter: ET470/40x, Emitter: ET525/50m, Dichroic: T495LP) for GFP-expressing strains. Fluorescence and transmitted light images were taken with a Photometrics CoolSNAP HQ^2^ cooled CCD camera, interfaced with LabView through the Bruxton Inc. SIDX API. We used a 10x objective, with a numerical aperture of 0.30, because it provided a wide field of view while allowing easy observation of individual cells.

Image acquisition was done with an in-house LabVIEW program, incorporating autofocusing in bright field followed by fluorescence imaging with automatically adjusted exposure time to avoid camera saturation. Optimal exposure times for fluorescence imaging may vary (~0.1-1 second). When we imaged *ade8*- cells using a particular light configuration, the very short exposure time (0.05 sec for initial images) created problems for image analysis. This could be due to the latency in shutter opening/closing becoming important in short exposure times. Alternatively, since high-intensity light was used (and hence the short exposure time), exposure time had to be reduced as cells grew to avoid camera saturation, and adjusting for variable exposure time during data analysis could introduce errors. When we added a neutral density filter and/or adjusted the size of light aperture so that the exposure time was ~0.3 sec, data analysis became normal. Four locations per well were imaged, with ~20-200 initial cells per image.

### Lid warmer

During extended imaging, condensation could accumulate on the underside of microplate lid, even in the temperature-controlled chamber. We encountered failures in autofocusing due to heavy condensation, and condensation can degrade proper Köhlier illumination. In order to prevent condensation, we used an optically transparent heating plate to warm the lid (Figure 1B), which kept the lid temperature an average of 0.68°C with 2σ of 0.22°C higher than the stage temperature (Supp Fig 1). This eliminated condensation (Figure 1C).

Our lid warmer used an ITO glass heating plate, with an integrated thermistor temperature sensor (Oven Industries TR91-170). The 1 mm-thick ITO glass had a 140nm ITO coating (from SPI Supplies), transmitted 88% of visible light, and had a sheet resistance between 30 - 60 Ohms/sq. We cut a 70mm x 120mm sheet with beveled corners to match the microplate lids. To apply power for heating, we affixed strips of 5 mm-wide conductive copper foil tape to the ITO coating along two opposite ends of the heating plate, and applied silver conductive epoxy (MG Chemicals, 8332-13G) to the tape, extending 1-2 mm onto the ITO coating to ensure a reliable connection. Wires were connected through brass conductors, which were insulated with heat shrink tubing and epoxied to the edge of the heating plate. We measured the resistance of the plate to be 20 Ohms, so the application of 5V DC generated (5 V)^2^/26 Ohms = 1.0 W. This provided the needed heat when applied. The sensor readings were tested, and adjusted if necessary, using a Barnant 115 thermocouple thermometer with a T type probe.

The temperature of the heating plate was controlled by a LabVIEW program, with the aid of a DAQ (National Instruments USB-6008) for reading temperature sensors. Based on the measured temperatures of the heating plate and the microscope stage, the LabVIEW program activated or deactivated the heating plate when the temperature difference was ≤0.5°C or ≥0.8°C, respectively. As discussed above, the heating plate was run at a fixed power of ~ 1W when active. Note: The 5V supply from the DAQ was used for the sensors, and a separate supply was used for heating the plate.

### Setting up samples in microtiter plates

We used flat-bottom transparent microplates with wells joined together at the bottom by a continuous sheet of plastic, such as Costar 3370 96-well plates (rather than 96-well plates where the spaces between the wells was open to the surrounding environment and thus more susceptible to temperature fluctuations). In these plates, air warmed by the lid warmer thermally insulates the sides of individual wells, which improves temperature uniformity.

We filled the outermost wells of a 96-well microtiter plate with water to reduce evaporation, leaving up to 60 wells for imaging. We diluted cells to low densities (1000~5000 cells inoculated in 300 μl medium) to maximize our growth window and to minimize metabolite depletion during measurements. When assaying death in the absence of supplements, we added 2~10 fold more cells since there was little cell birth. We spun the plates at 2000 rpm for 2 minutes to settle all cells to the bottom of the wells, set up the microscope as described above, and imaged the same four positions in each selected well periodically (every 0.5~2 hrs). For each position, a bright field and a fluorescent image was saved. For growth assays, we ran experiments until fluorescence leveled off (16+ hrs for *lys2*-, and 30+ hrs for *ade8*- cells). For death assays, we ran experiments for up to 66 hr. We found that growth rates for the same samples did not vary significantly across different well positions in a plate (Supp Fig 10).

### Autofocusing

Automated imaging of the cells was performed using custom software written in LabVIEW. At each time point, it was necessary to auto-focus on the cells due to the small vertical drift caused by small changes in temperature or mechanical stress. At the beginning of an experiment, manual focusing was performed on four corner sample wells to ensure that the plate was level (otherwise, we need to adjust the screw positions of the plate holder). Then, coarse auto-focusing was performed for one position in each sample well at +/− 100 z-positions spaced at 3 μm apart, and the best focal plane was chosen to initiate an experiment. To identify the best focal plane for imaging and to prevent a loss of focus during the experiment, fine auto-focusing (+/− 30 z-positions spaced at 2 μm apart) was performed for each time point.

Each 16-bit image was imported directly into Labview, and converted into a twodimensional array of real numbers. The optimal z-position for focusing was chosen using a variant of the Brenner auto-focus algorithm ^20,21^. The quality of focus, *A*(*z*), was measured by computing the averaged horizontal and vertical gradient in each image:

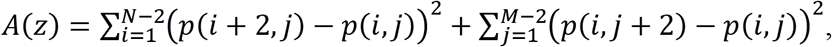

where *N* ⨯ *M* were the dimensions of each image, and *p*(*i,j*) was the pixel intensity at row *i* and column *j*. Local maxima in *A*(*z*) corresponded to sharply-focused images. Comparison between data two pixels apart rather than adjacent pixels reduced the effects of correlated noise and the natural illumination of nearby pixels.

When observing yeast we often observed multiple local maxima in the range of z positions, which was due to the focusing of light within the cells. Assuming yeast cells behave like small spherical lenses of diameter *D* ≈ 5 *μm*, then this phenomenon is likely a complex function of reflection, refraction, and diffraction ^23^. However, some qualitative features may be illustrated by simply considering refraction of a ball lens, where the effective focal length (*EFL*) is ^23^:

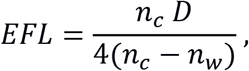

where *n*_*w*_ = 1.33 is the index of refraction of the surrounding water, and *n_c_* is the index of refraction of the cell. Supp Fig 12 shows that the first maxima is due to light that is focused approximately 14 μm from the yeast cells, which would correspond to *n*_*c*_ = 1.46. This is consistent with previous measurements showing that *n_c_* ≈ 1.53 ^23^, although we note that this value depends on the size and density of the cells. The second maxima is located at an image plane below the cells that contains light halos from the apparent source of the focused light.

We found that the minimum in *A*(*z*) between these two maxima conveniently corresponded to a focal plane adequate for identifying and imaging the cells (Supp Fig 12). Thus, we choose this local minimum for imaging in all of our experiments. The three points closest to the minimum are fit to a parabola, and the minimum of the parabola is chosen as the optimal focus position. The microscope stage is then moved to the optimal focal plane for imaging. A similar method for autofocusing, i.e. using the local minimum in the autofocus score, has been used before ^24^.

### Image analysis

We analyzed time-lapse images using Bioact2, an ImageJ plugin written by Adam Waite (available at https://github.com/nodice73/Java/tree/master/imagej_plugins/bioact). Bioact2 measured the background-subtracted total fluorescence intensity of all cells in an image. To distinguish fluorescent cells from background, each fluorescent image was blurred with a Gaussian filter using a standard deviation (σ) of 1 pixel. Low frequency noise was removed using the “rolling ball” background subtraction algorithm ^25^. The dynamic range of the image was reduced from 16-bit to 8-bit, and each pixel was replaced by the maximum value in a 3 pixel radius. The resulting image was thresholded using the “maximum entropy” method for low density images or the “iso data” method for higher-density images and converted to a binary mask ^26^. After filling holes using binary closing, the mask was dilated by 2 pixels. From this mask, the percent pixels considered foreground was calculated. Each mask was then applied to the un-manipulated original image, and the foreground and background intensities were measured. At low cell densities, the background was calculated for each image and subtracted. We found that when the foreground made up more than a specified fraction of the total image area, the background estimate was no longer accurate, and the running average background value calculated before this threshold was met was used as the background value for subsequent images. This method occasionally failed if cells were nearly confluent during late stage growth. However, we were only interested in maximum growth rate, which occurred before confluency.

We plotted background-subtracted fluorescence intensity over time for all four positions in each well to allow visual inspection. In rare occasions, all four positions were out-of-focus and none were used. In a small subset of experiments, a discontinuous jump in data appeared in all four positions for reasons we do not understand. We did not calculate rates across the jump.

Occasionally, data from one or two positions deviated from the rest. This could have been due to a number of reasons, including spurious shifts in stage position, or black or bright dust particles in the field of view. In these cases, we inspected the images and outliers with obvious causes were excluded. If the fluorescence dynamics of four positions differed due to cell heterogeneity at low concentrations of metabolites, all positions were retained.

We normalized intensity against that of time zero, and averaged across positions. We calculated growth rate over three to four consecutive time points, and plotted the maximal net growth rate against metabolite concentration. If maximal growth rate occurred at the end of an experiment, then the experimental duration was too short and data were not used.

### Individual cell tracking at low metabolite concentrations

At very low concentrations of supplements, we manually counted birth and death events by scanning through fluorescent images in ImageJ (e.g. Supp Movie 2-3). After counting the initial number of cells in an image, we proceeded through each image and noted the appearance or disappearance of cells. We counted a birth event as the appearance of a new cell adjacent to a cell present in the previous image. We differentiated this from two vertically-stacked cells shifting orientation to create the appearance of a new cell birth, as the fluorescence intensity of stacked cells dropped noticeably during orientation shift. We counted a death event when a cell present in the previous image suddenly lost fluorescence. Occasionally cellular fluorescence slowly faded over time rather than disappearing suddenly, and in this case we counted the initial drop in intensity as the death of the cell.

### Bioassay quantification of metabolite concentrations

We used a bioassay to quantify lysine concentrations from supernatants. In order to obtain supernatant, we filtered cell cultures through a 0.45 μm nitrocellulose filter and stored at -80°C until quantification. We mixed 150 μl sample with an equal volume of master mix containing 2x SD and WY2270 lysine-requiring tester cells (~1×10^4^ cells/ml) in a flat-bottom 96-well plate.

We measured growth rates of WY2270 in unknown samples and compared them to growth rates of WY2270 run concurrently with known concentrations of lysine. The growth rate scaled linearly with lysine concentration up to 1 μM (i.e. 2 μM in undiluted sample, Supp Fig 11). From this standard curve, we inferred lysine concentrations of samples. Assay sensitivity was 0.1 μM.

### Quantifying population dynamics using flow cytometry

We first prepared bead standards for quantifying cell density. Fluorescent beads (ThermoFisher Cat R0300, 3 μm red fluorescent beads) were autoclaved in a factory-clean glass tube, diluted into sterile 0.9% NaCl, and supplemented with sterile-filtered Triton X-100 to a final 0.05% (to prevent beads from clumping). We sonicated beads and kept them in constant rotation to prevent settling. We quantified bead concentrations by counting beads using a hemacytometer and a BD Coulter counter. The final bead stock was generally 4~8×10^6^/ml.

Culture samples were diluted to OD 0.01~0.1 (7×10^5^~7×10^6^/ml) in MilliQ H_2_O in unautoclaved 1.6ml Eppendorf tubes. In a 96-well plate, 90 μl sample were supplemented with 10 μl bead stock to calculate cell density from cell:bead ratio. We also added 2 μl of 1 μM nucleic acid dye ToPro3 (Molecular Probes T-3605) which stains dying/dead cells with compromised membrane. Flow cytometry was performed on Cytek DxP Cytometer equipped with four lasers, ten detectors, and an autosampler. Fluorescent tags GFP, mCherry, and ToPro were respectively detected by 50 mW 488 nm laser with 505/10 (i.e. 500~515nm) detector, 75 mW 561nm Laser with 615/25 detector, and 25mW 637nm laser with 660/20 detector. Each sample was run in triplicate and individually analyzed using FlowJo^®^ software to identify numbers of events of beads, dead cells, and various live fluorescent cells. We calculated the mean cell density from triplicate measurements, with the coefficient of variation generally within 5%~10%.

### Chemostat culturing

We constructed an eight-vessel chemostat with a design ^13^ modified from an existing multiplexed culturing device ^27^. A chemostat creates a nutrient-limited environment where the population is forced to grow at a constant, pre-determined rate slower than the maximal growth rate ^28^.

Specifically, a medium containing a limiting metabolite is added to the culturing chamber at a constant flow rate *f*(ml/h). The culture effluent is removed from the chamber at the same rate *f*, thereby maintaining a constant culture volume *V*. Mathematically ^28^, live population density *N* satisfies

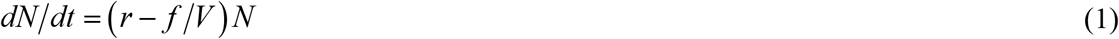

where *r* (h^−1^) is the net growth rate (birth rate minus death rate), *f* is the flow rate, and *f/V* is the dilution rate (h^−1^). At steady state, the net growth rate and the dilution rate are equal:

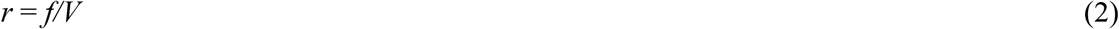

The limiting metabolite also reaches a steady state in the culturing vessel. Thus, the steady state concentration of the limiting metabolite supports a net growth rate equaling the dilution rate.

Due to rapid evolution, we designed experiments so that live and dead populations quickly reached steady state and the experiment lasted ≤26 hours (bioRxiv). We washed exponentially-growing cells to remove extracellular lysine and inoculated 1/4~1/2 of the volume at 1/3 of the expected steady-state density. We filled the rest of the 19ml chamber with reservoir media (resulting in less than the full 20 μM of reservoir lysine, but more than enough for maximal initial growth rate, ~10-15 μM). We sampled cultures periodically to track population dynamics using flow cytometry (Supp Fig 7A) and filtered supernatant through a 0.45 μm nitrocellulose filter. We froze the supernatants for metabolite quantification at the conclusion of an experiment (Supp Fig 7B).

## Acknowledgement

We thank Jose Pineda for performing the experiment in Supp Fig 2

---

**Supp Fig 1.**
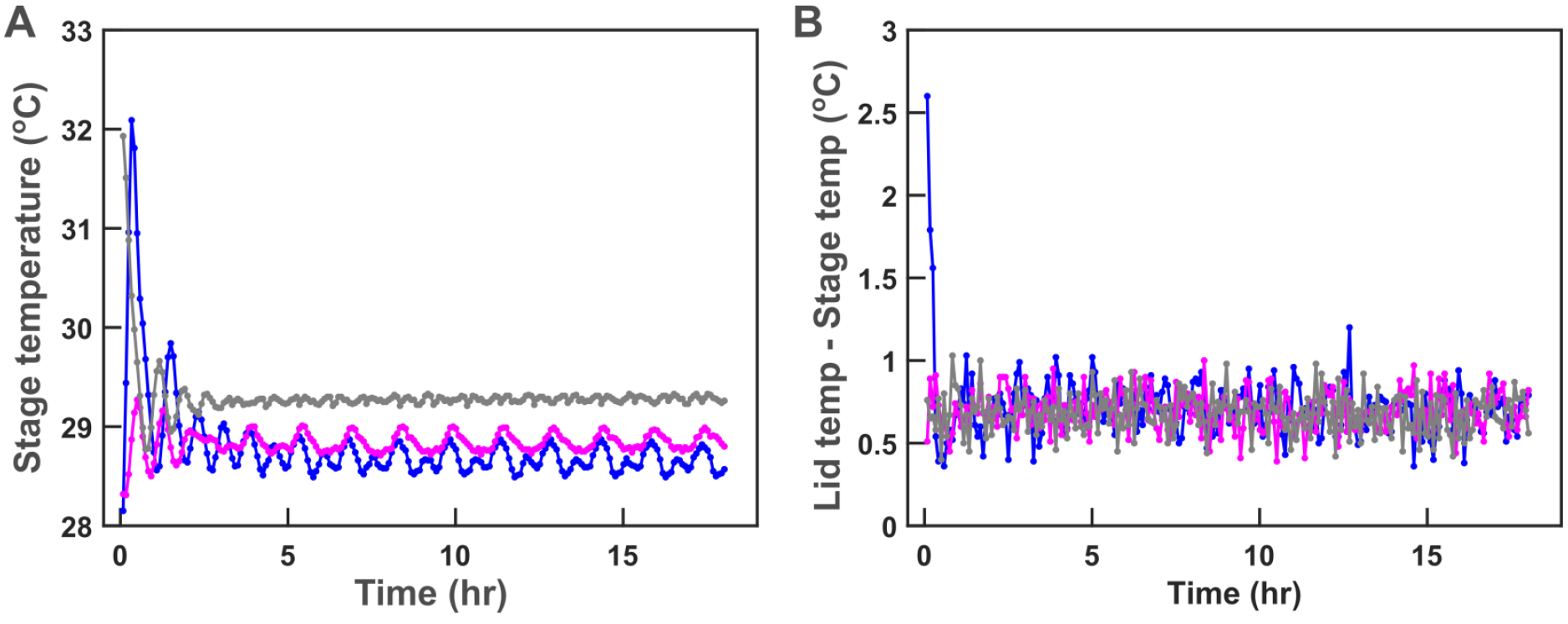
Temperature-controlled stage and lid warmer. **(A)** Stage temperature was set to 29.5 °C and was maintained at within 0.5°C of the set temperature except for the initial period of time. **(B)** The lid warmer maintained the microplate lid 0.5~ 0.8°C above the stage temperature. The average temperature difference was 0.68°C. A LabVIEW program compared temperature readings from the stage and from the lid warmer, and turned on and off the heater to maintain the set range of temperature difference. The initial temperature overshoot may be due to non-equilibrium distribution of heat in the system. Three colors indicate three independent experiments. Growth rates at 5 μM (as seen in Figure 4) for all three experiments were similar (0.50 +/− 0.02/hr) despite the observed ~0.5°C differences in temperature.

**Supp Fig 2.**
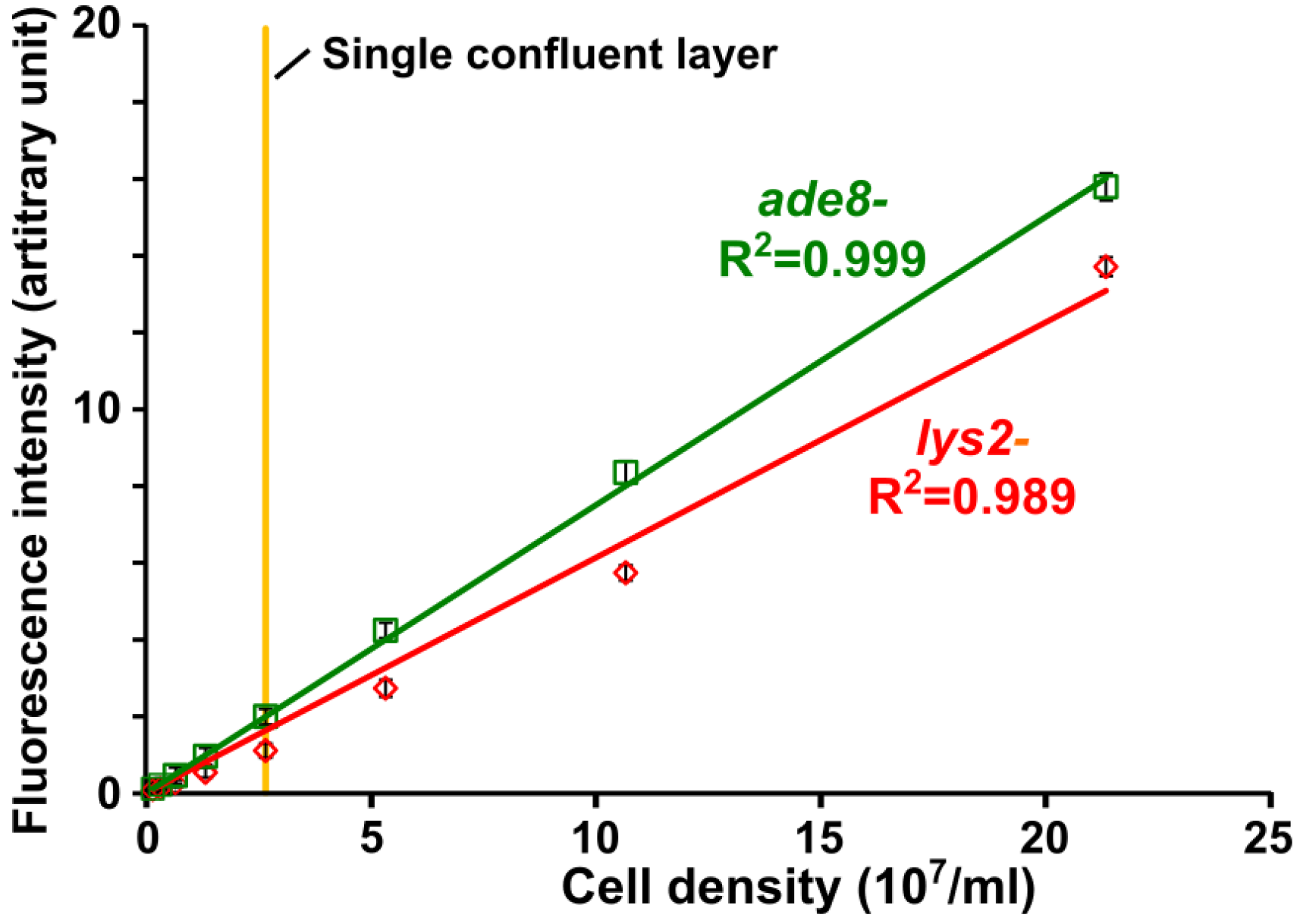
A linear relationship between total live cell number and total fluorescence intensity in a field of view. Various numbers of live cells (counted by flow cytometry) were placed in minimal medium lacking glucose (to arrest cell growth) in microtiter wells and allowed to settle to the bottom of wells. The plate was then imaged using our microscope setup to measure the total fluorescence intensity. For both red (mCherry-expressing *lys2*-) and green (GFP-expressing *ade8*-) fluorescent strains, total cell numbers and total background-subtracted fluorescence intensities in a field of view display linear relationships. At the highest density, there were at least nine cell layers.

**Supp Fig 3.**
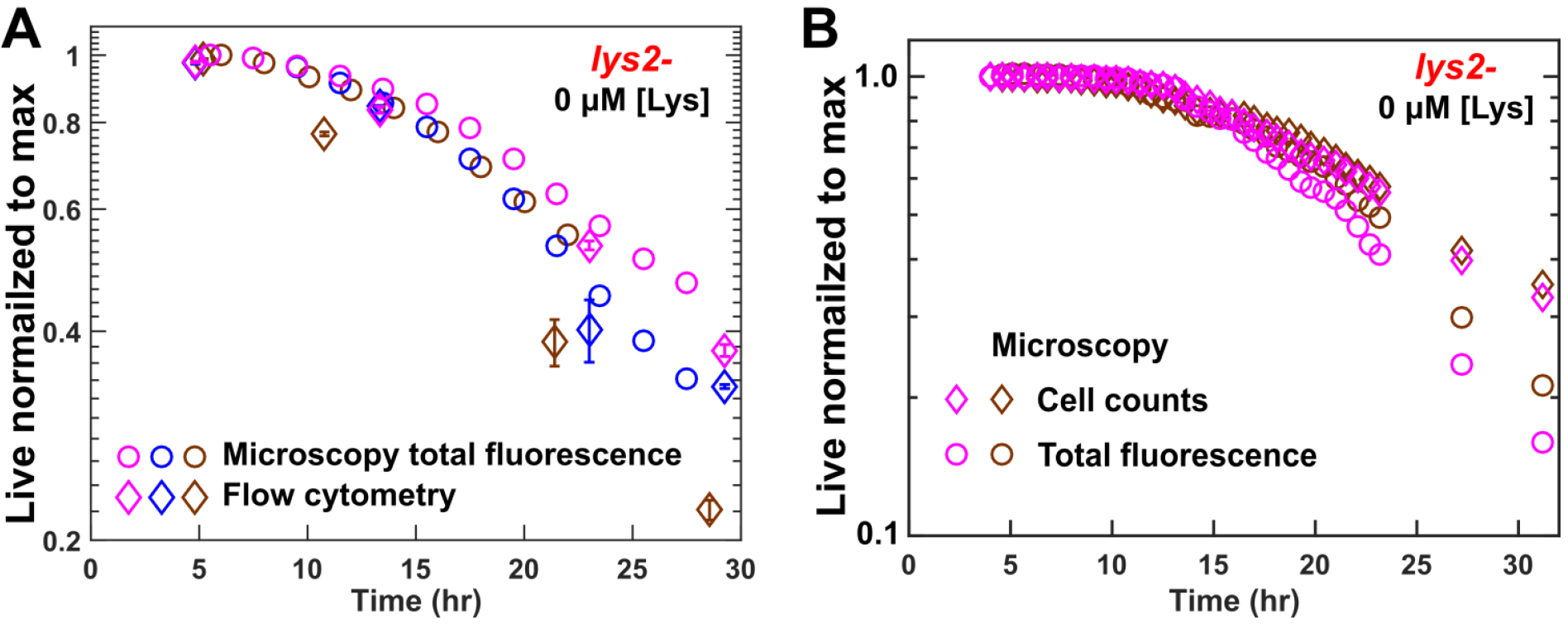
Microscopy and flow cytometry measurements reveal multi-phasic death dynamics of lys2- cells. We grew *lys2*- (WY1335) in excess lysine. At time zero, we washed cells free of lysine, and prestarved cells for 4~5 hrs to deplete vacuolar lysine storage. We then diluted cells in SD (<20,000 cell/ml) and imaged them. **(A)** Microscopy total fluorescence intensity (circles) and flow cytometry (diamonds) revealed multi-phasic death kinetics (slow death rate followed by faster death rate). For a fraction of wells, we also periodically harvested and concentrated cells for flow cytometry measurements. Error bars are 2x standard deviations from multiple independent measurements (three technical replicates for FACS, and tens of images for fluorescence imaging). **(B)** Cell counts (diamonds) and total fluorescence intensity (circles) reveal multi-phasic death kinetics. In cell counting, ~200 cells were followed. In each panel, different colors represent experiments done on different days.

**Supp Fig 4.**
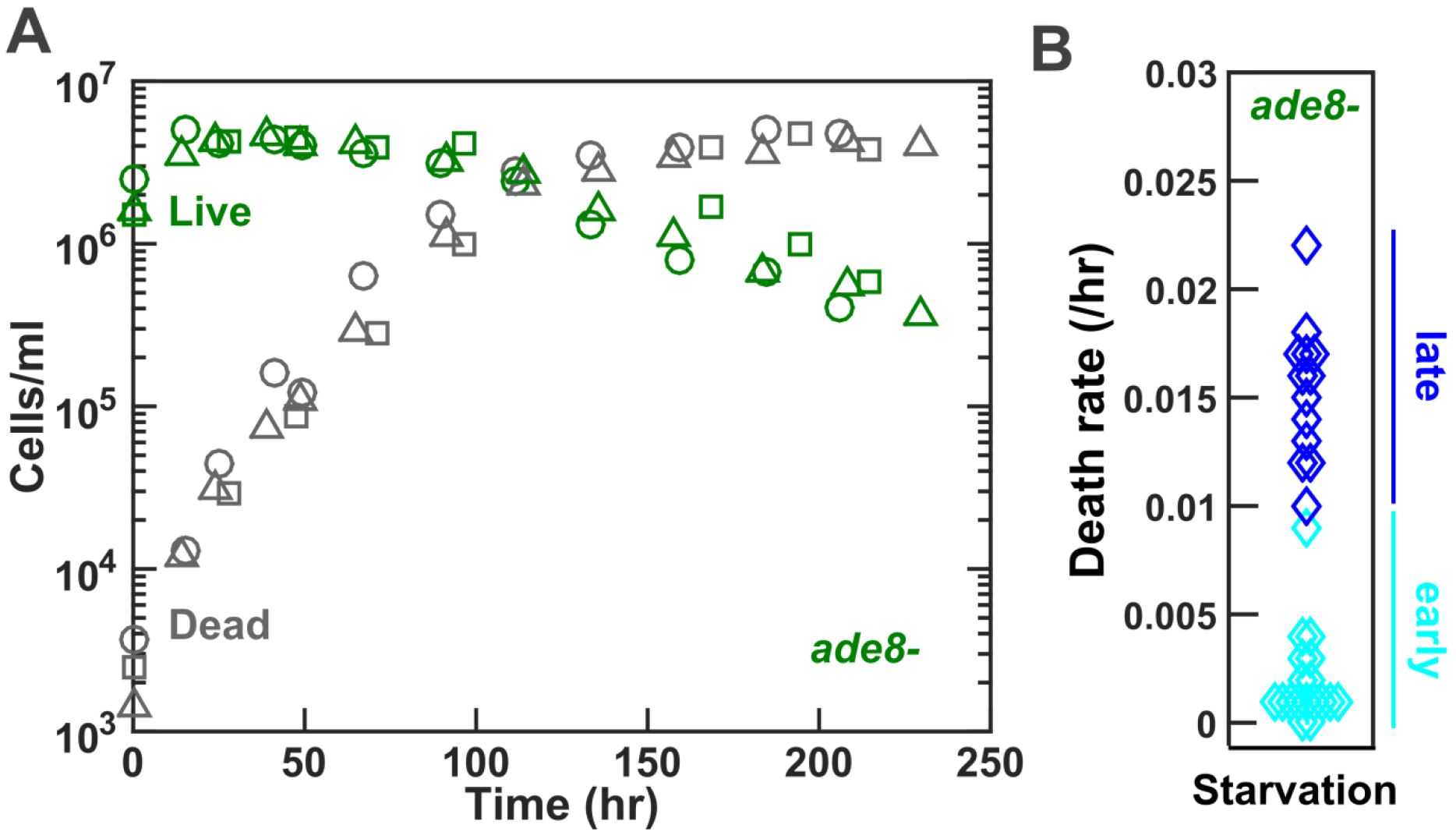
Time-dependent death rate of ade8- during starvation. **(A)** Death dynamics of *ade8*- cells. Exponential *ade8*- (WY1340) cells were washed and starved. We tracked dead and live cell densities over time using flow cytometry. **(B)** Death rate is initially slow and then speeds up. Early death rates were generally calculated within 24-96 hr, while late death rates were generally calculated within 70-230 hr.

**Supp Fig 5.**
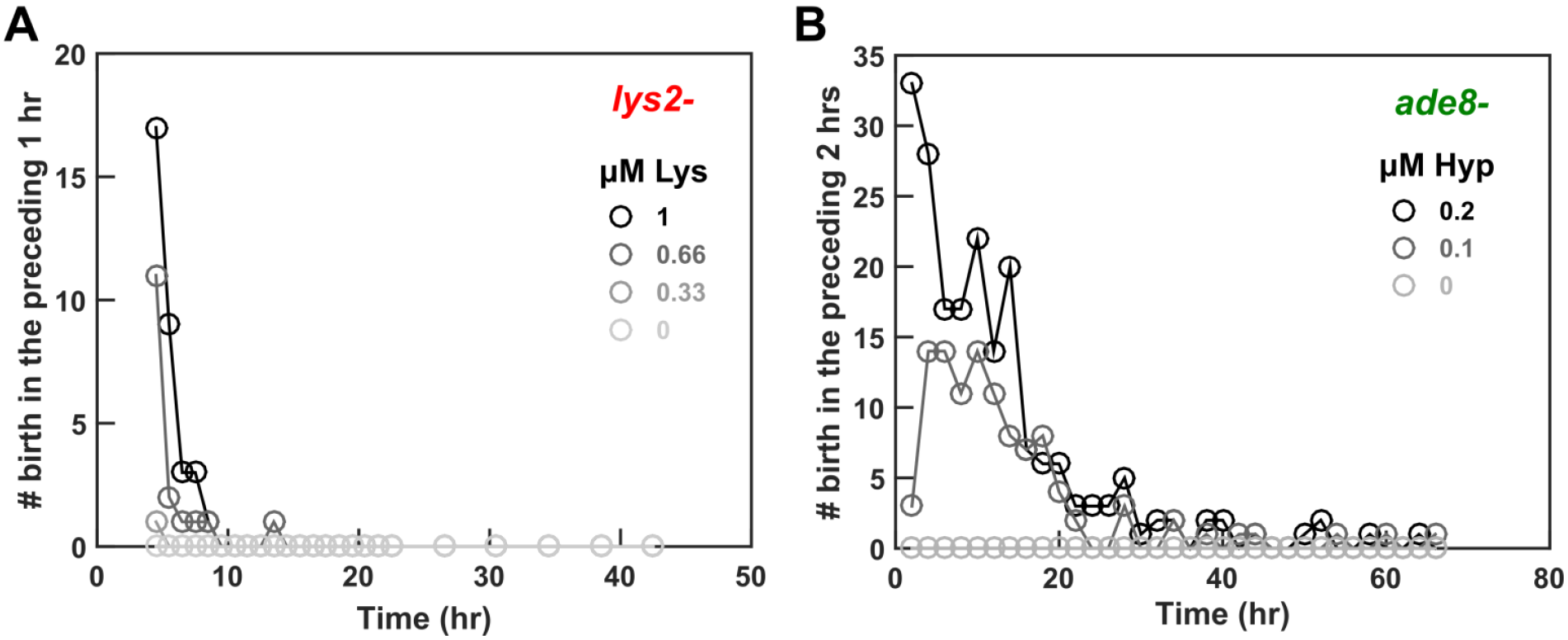
Low concentrations of the required metabolite supported birth for ade8- but not lys2- cells. **(A)** *lys2*- cells grown in excess lysine were washed free of lysine at time zero, and the number of birth over a 1-hr interval was counted. Data source was identical to that in Figure 2A. (B) We grew *ade8*- in excess hypoxanthine to exponential phase, washed away hypoxanthine, and prestarved cells for 24 hrs. At time zero, we started imaging. The number of birth over a 2-hr interval was counted. Data source was identical to that in Figure 3D.

**Supp Fig 6.**
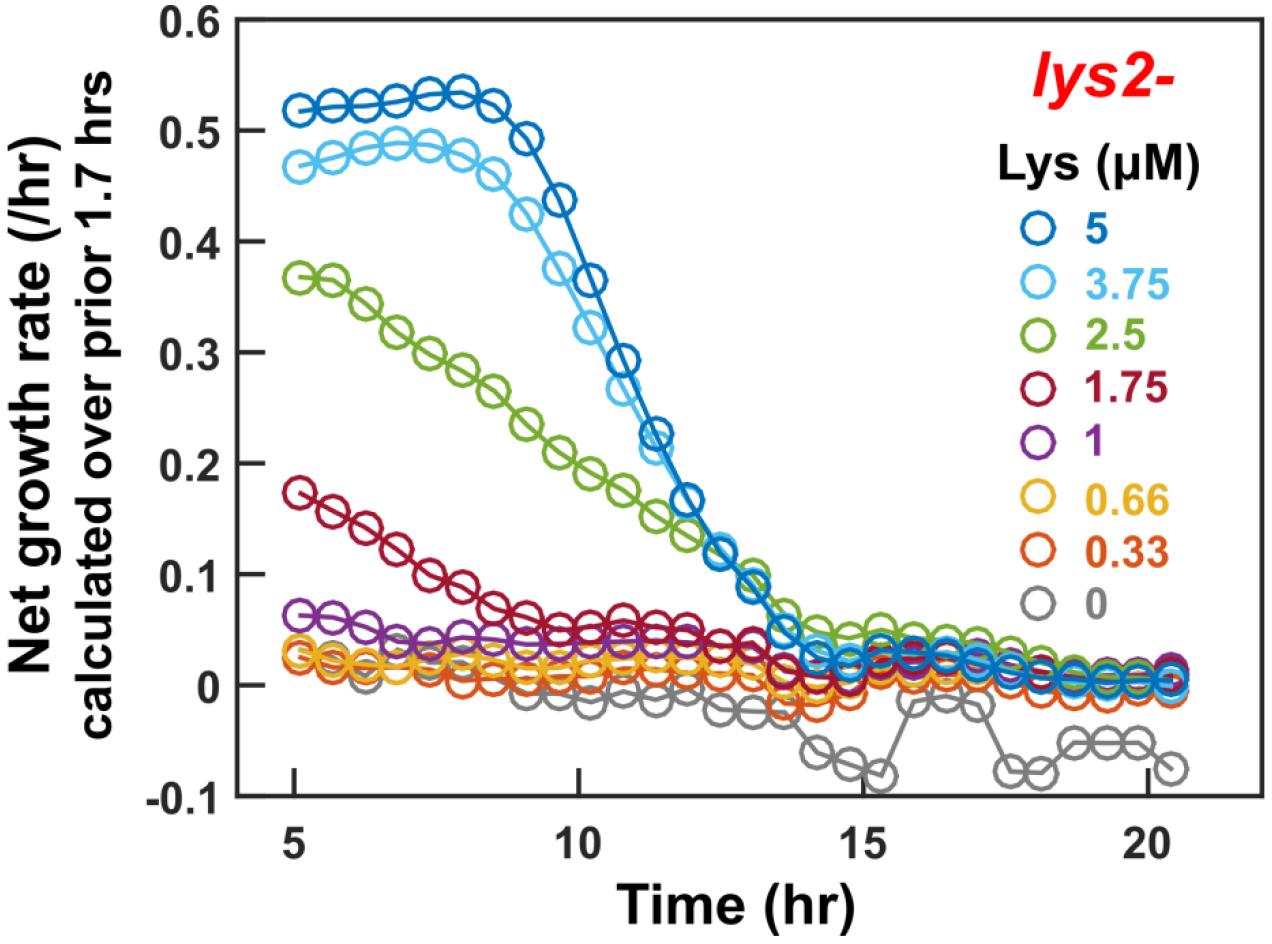
Steady growth rate is maintained only for ≥2.5 μM lysine. We calculated growth rates from fluorescence intensity over time. Each data point corresponds to the growth rate calculated over the four previous time points, a time window of approximately 1.7 hours. For ≥2.5 μM lysine, cells reached a maximal growth rate and maintained it across at least two sliding windows (e.g. hours ~5-8 for 3.75 and 5 μM). For 1.75 μM lysine, the growth rate declined throughout the duration of measurements. For ≤1 μM lysine, any positive net growth rate after residual growth was due to cell swelling not cell birth (Figure 3C).

**Supp Fig 7.**
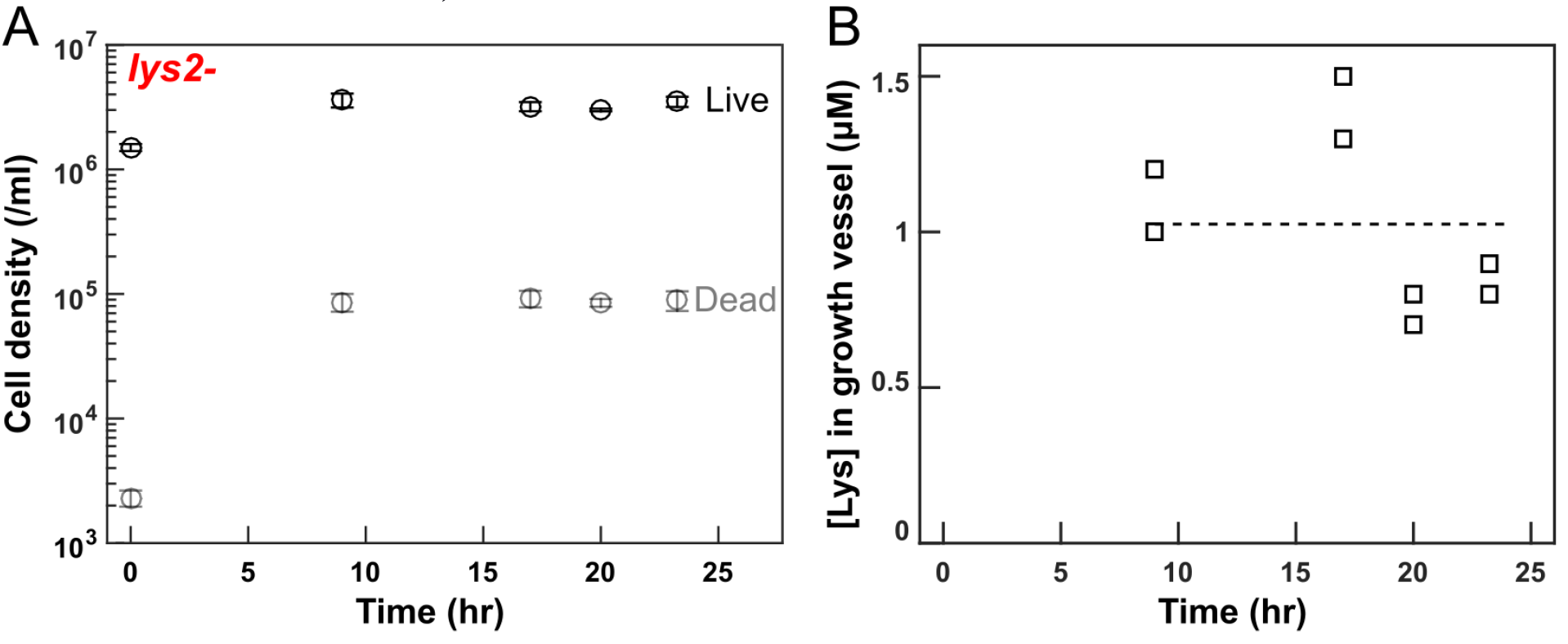
Chemostat dynamics rapidly reach a steady state. *lys2*- cells growing exponentially in excess lysine were washed free of lysine, and inoculated in a chemostat vessel at 1/3 steady-state density in the presence of ~10-15 μM lysine. Minimal medium containing 20 μM lysine was dripped into the culturing vessel (19 ml) at a set rate to achieve desired growth rate (8-hr doubling time corresponding to a flow rate=19 ml*ln(2)/8 hr = 1.646 ml/hr; “Chemostat culturing” in Methods). We tracked live and dead cell densities **(A)** using flow cytometry (Methods, “Quantifying population dynamics using flow cytometry”) and tracked lysine concentrations **(B)** using a bioassay (Methods, “Bioassay quantification of metabolite concentrations”).

**Supp Fig 8.**
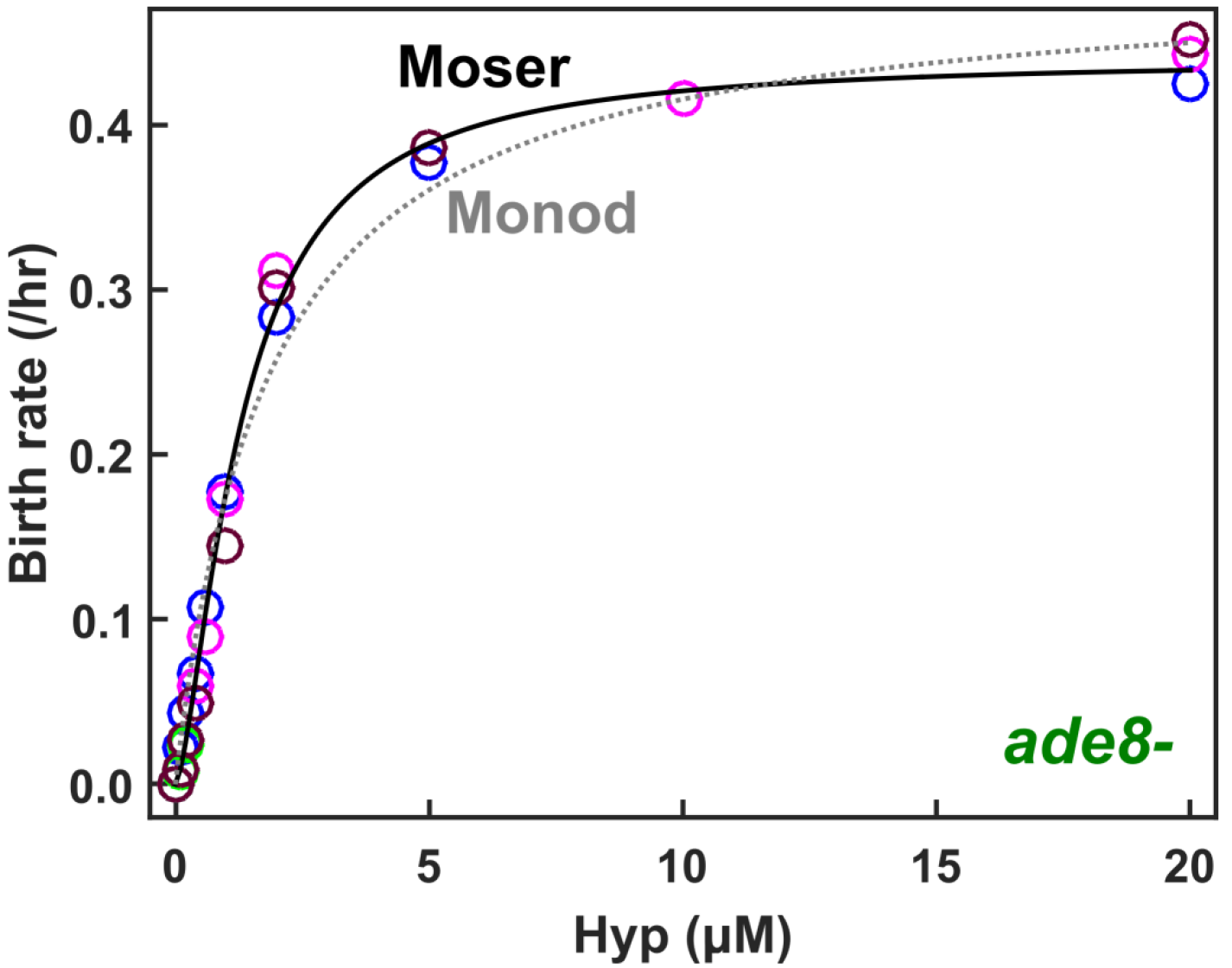
Birth rates of ade8- as a function of hypoxanthine concentrations. *ade8*- (WY1340) cells were grown to exponential phase in minimal medium supplemented with excess hypoxanthine, washed free of hypoxanthine, and pre-starved for 24 hrs to deplete cellular storage. Cells were then incubated at various concentrations of hypoxanthine and imaged. The Monod model (grey dotted line) predicted a maximal birth rate of 0.49/hr (95% CI: 0.46~ 0.52/hr). In the Moser model (black), maximal birth rate *b*_max_ = 0.44/hr (95% CI: 0.43 ~ 0.45), hypoxanthine concentration for half maximal birth rate *K*_*m*_ = 1.3 μM (95% CI: 1.2 ~ 1.4), and *n* =1.5 (95% CI: 1.4~ 1.7). In comparison, experimentally-measured *b_max_* is 0.44±0.03/hr.

**Supp Fig 9.**
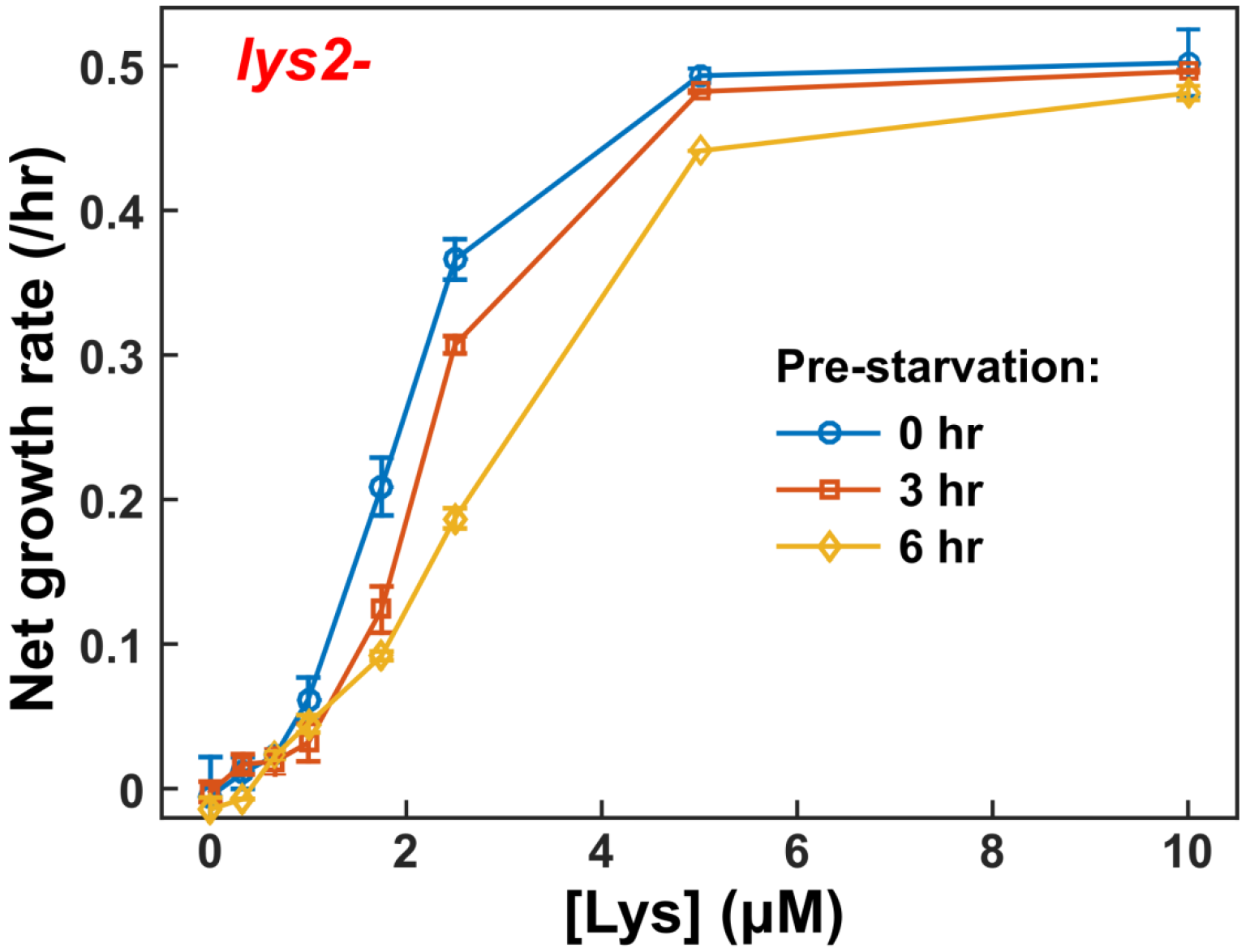
The duration of pre-starvation can affect growth rates. Pre-starvation at zero lysine, which could be used to deplete cellular lysine storage, reduces net growth rate compared to no pre-starvation. *lys2*- cells were grown to exponential phase in excess lysine. At time zero, we washed cells free of lysine, and pre-starved them for 0, 3, or 6 hrs before imaging them in various concentrations of lysine. We analyzed data from hour 3 and on (post-residual growth). Using 3-hr sliding time windows, we calculated the net growth rate over time. We plotted maximal growth rate with error bars indicating 2x error of slope estimation. Note that we used fluorescence to quantify growth rate and did not correct for the absence of birth in 0~1μM lysine. This resulted in perceived positive net growth rates.

**Supp Fig 10.**
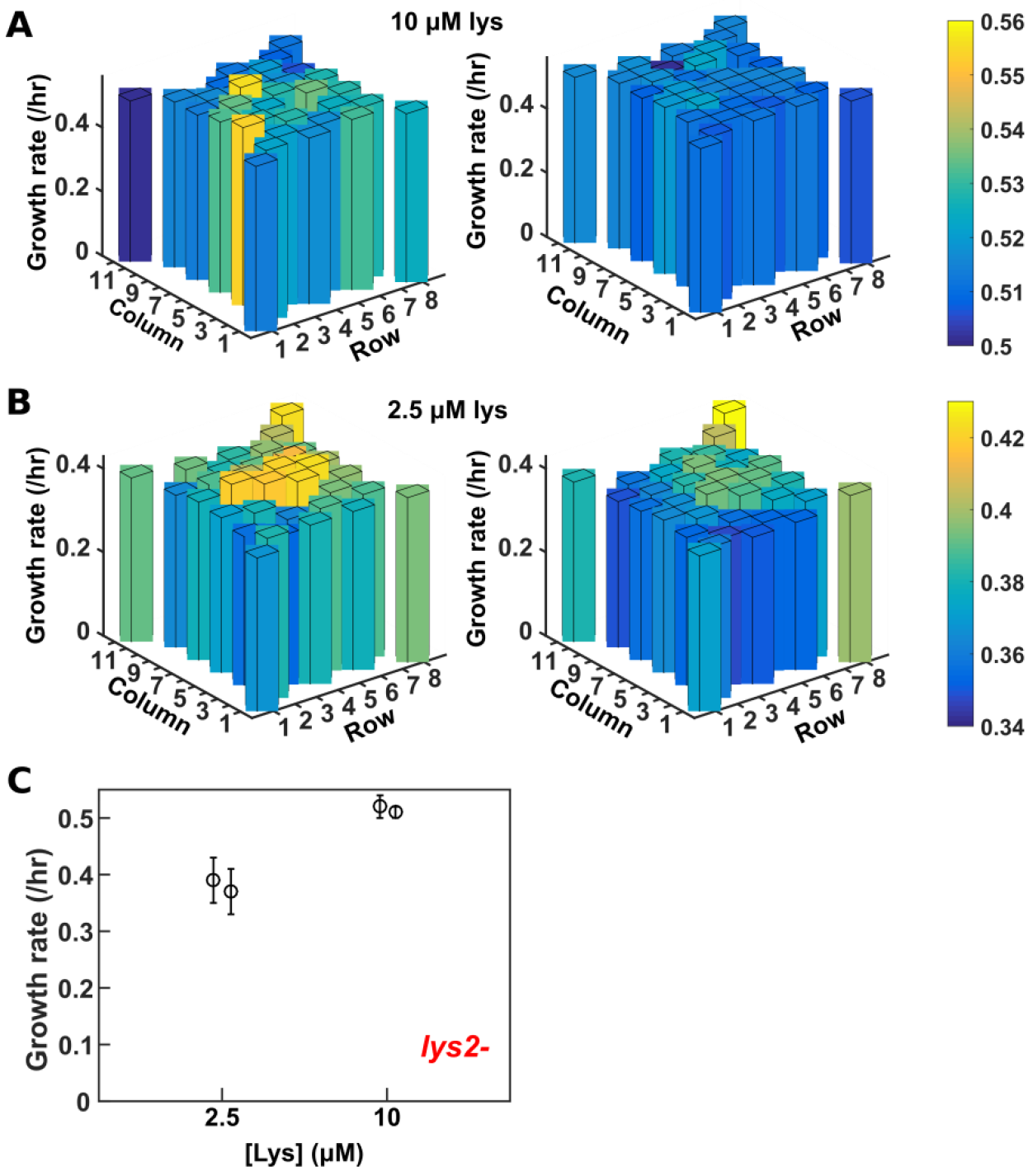
Well location does not significantly impact growth rate. To examine potential well-position bias in growth rate, we quantified growth rates of *lys2*- cells across various wells of a 96-well plate at an identical initial lysine concentration. **(A)** We did not observe any correlation between well position and growth rate at 10 μM (non-limiting) lysine in either of the two experiments (left and right). **(B)** There was a slight correlation between well position and growth rate at 2.5 μM (limiting) lysine. Wells that supported faster growth in one experiment tended to support faster growth in the other (left vs right). This is likely the result of the order in which cells were imaged, as imaging could take over an hour per round, and at 2.5 μM lysine, growth rate continuously declined during measurement (Supp Fig 6). **(C)** Growth rates averaged across wells in a plate were similar between two independent experiments. Error bars represent 2*standard deviations.

**Supp Fig 11.**
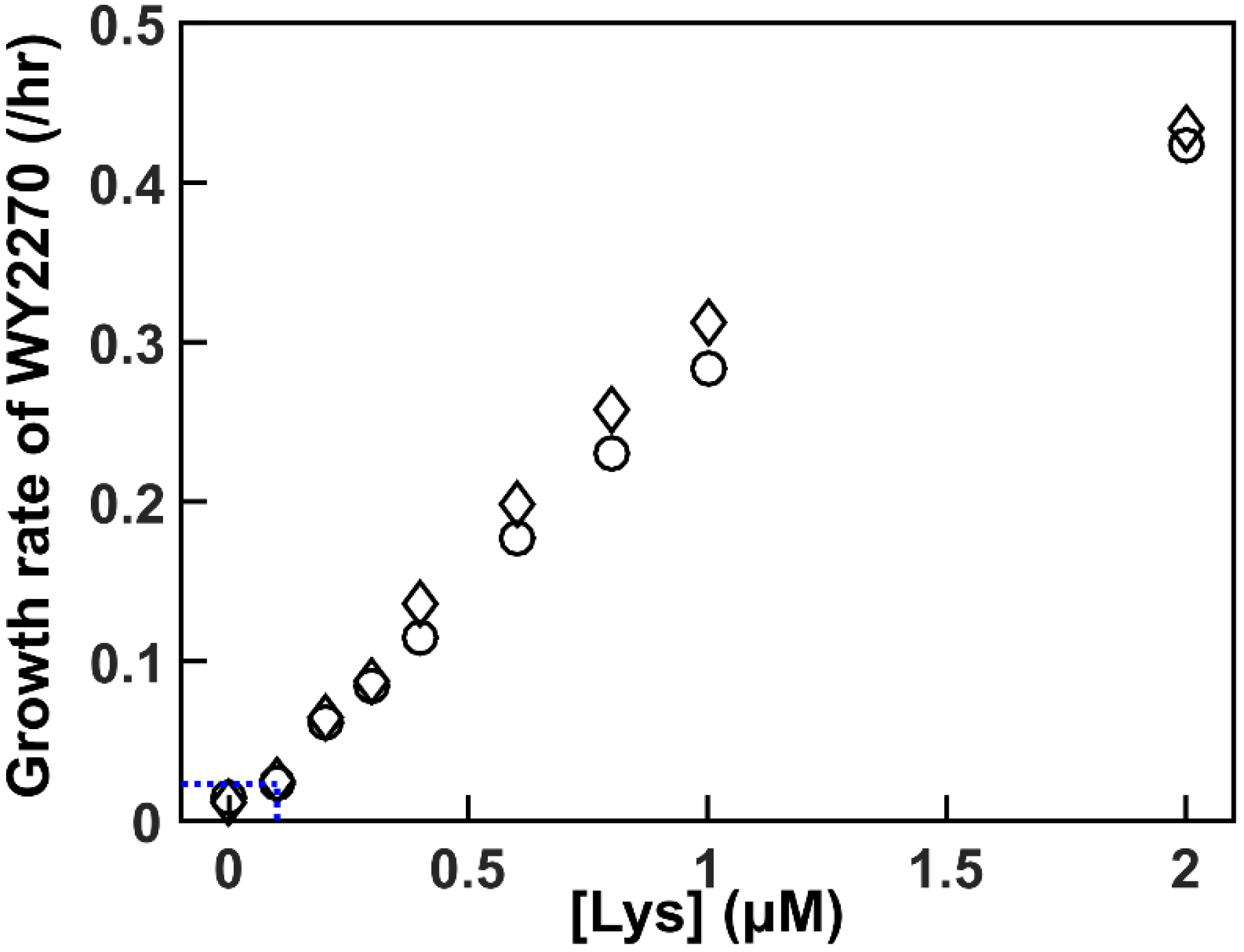
Using an evolved clone to measure low concentrations of lysine. WY2270, an evolved *lys2*- clone with significantly improved affinity for lysine compared to WY1335, can detect lysine as low as 0.1 μM. Dotted line marks detection limit. Circles and diamonds mark two independent replicates.

**Supp Fig 12.**
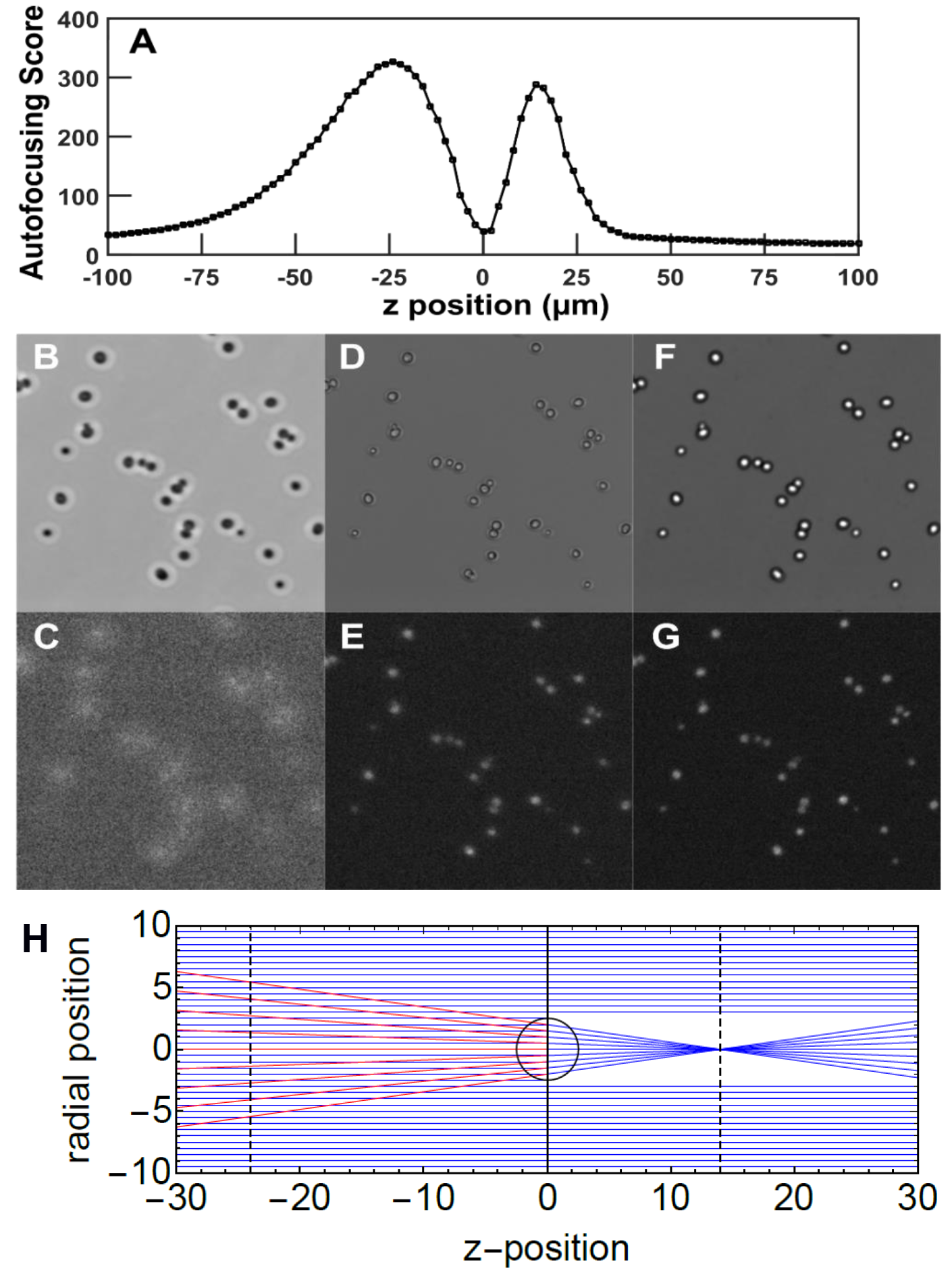
Autofocusing. Adequate fluorescent images can be obtained by choosing the local minimum of the autofocus score as our imaging plane. **A)** The autofocus score *A(z)* had two local maxima, corresponding to image artifacts created by lensing effects of yeast cells. Bright field **(B-F)** and fluorescent **(C-G)** images were taken near z = −24 μm, 0 μm, and 14 μm, corresponding to local maximal, minimal, and maximal *A(z)*, respectively. The yeast cells were in focus at the local minimum. **(H)** A qualitative explanation of autofocusing. This figure shows the apparent origin of light rays reaching the camera. The density of light rays in a given focal plane corresponds to the apparent brightness. The optical effects are due to focusing of the light rays (blue) by the yeast cells (black circle). The image plane at z = 14 μm (dashed line) contained a sharp bright spot (which looks bigger than a point due to diffraction), and the image plane at z = −24 μm (dashed black line) contained bright halos due to the apparent source of the focused light rays (red lines).

Supp Movie 1 Time-lapse fluorescence microscopy of lys2- at high lysine

Supp Movie 2 Time-lapse fluorescence microscopy of lys2- at low (0.66 μM) lysine

Supp Movie 3 Time-lapse fluorescence microscopy of lys2- at zero lysine

Supp Movie 4 ade8- cell re-dividing after having transiently lost fluorescence.

Supp Movie 5 Time-lapse bright field microscopy of lys2- at zero lysine

